# Multimodal analysis of *in vitro* hematopoiesis reveals blood cell-specific genetic impacts on complex disease traits

**DOI:** 10.1101/2025.05.23.655825

**Authors:** Rong Qiu, Khanh B. Trang, Carson Shalaby, James A. Pippin, James Garifallou, Struan F.A. Grant, Christopher S. Thom

## Abstract

*In vitro* hematopoiesis systems can be used to define mechanisms for blood cell formation and function, produce cell therapeutics, and model blood cell contributions to systemic disease.

Hematopoietic progenitor cell (HPC) production remains inefficient, precluded by knowledge gaps related to specification and morphogenesis of specialized hemogenic endothelial cells, which undergo an endothelial-to-hematopoietic transition (EHT) to form HPCs. We elected to define changes in gene expression and chromatin organization during HPC formation to reveal regulatory mechanisms. Using paired single cell RNA/ATAC sequencing together with Hi-C, we profiled cells before and after EHT. Pathway analysis and pseudotime inferences confirmed a continuum of stromal and endothelial cells undergoing development into HE cells and lineage-based HPCs *in vitro*. In these cell types, we characterize cis-regulatory elements and transcriptional regulatory activities that facilitate EHT and HPC homeostasis, including for SNAI1, SOX17, TGFβ, STAT4, as well as for GFI1b and KLF1 in megakaryocyte- and erythroid-biased progenitors, respectively. We then leveraged our insights into chromatin organization among *in vitro*-derived cells to assess enrichments corresponding to human trait variation reported in human genome wide association studies. HPCs revealed locus enrichment for quantitative blood traits and autoimmune disease predisposition, which were particularly enriched in myeloid- and lymphoid-biased populations. Stromal and endothelial cells from our *in vitro* cultures were specifically enriched for accessible chromatin at blood pressure loci. Our findings reveal genes and mechanisms governing *in vitro* hematopoietic development and blood cell-related disease pathology.

## Introduction

Hematopoietic progenitor cells (HPCs) are a lifelong source of circulating blood and immune cells.^1,2^ During embryonic life, HPCs arise from specialized hemogenic endothelial cells (HECs) through an endothelial-to-hematopoietic transition (EHT)^3–5^. EHT is an evolutionarily conserved process that can be recapitulated *in vitro* using human induced pluripotent stem cell (iPSC)-based culture systems.^6,7^ These *in vitro* systems offer an opportunity to dissect how developmental programs impact HPC formation and function. *In vitro*-derived HPCs and hematopoietic cells also offer an opportunity to produce cells, including rare cell types, in ample quantities to dissect how these cells contribute to complex disease pathology from a functional genomics perspective.

Current *in vitro* systems can produce engraftable stem cells and mature blood cells, but inefficient derivation of functional HPCs remains a major barrier for blood-based regenerative therapies.^8,9^ Addressing this limitation requires a detailed understanding of the molecular, epigenetic, and regulatory mechanisms that orchestrate early hematopoietic development. We and others have functionally validated loci and genes identified in human genome wide association studies (GWAS) using *in vitro* approaches^10–12^.

Gene predictions are largely based on locus proximity, given a paucity of chromatin-level data from *in vitro* systems. Thus, there remains a conceptual challenge in leveraging *in vivo* human genetic insights to optimize *in vitro* HPC production. Do *in vitro* models faithfully recapitulate the genomic regulatory landscape of human development? Evidence for this relationship could come from *in vitro*-derived blood cells being enriched for human GWAS loci. The *in vitro* system could then specify key developmental stage(s) wherein chromatin regulation is altered, implicated not only genomic loci but related genes and cell stages^13^. Although gene expression changes appear consistent across *in vitro* and *in vivo* hematopoiesis, there are not yet data to evaluate accessible chromatin changes during *in vitro* hematopoiesis, nor maps of the 3-dimensional chromatin looping events critical for gene regulation.

Current *in vitro*-derived blood cells can also define genetic mechanisms related to blood cell traits and blood cell-influenced disease susceptibility. Despite significant progress mapping genome-wide associations for hematologic trait and disease phenotype variations, a critical gap remains in understanding how variants functionally influence blood cell formation and function.^14–16^ Multimodal single cell profiling and functional genomics have provided insight into human disease trait variation using *in vitro* models^17^, including neurodevelopmental paradigms based on iPSC-derived cells^18^. Multimodal platforms based on analysis of *in vitro* and *in vivo* cells at single cell resolution have recently elucidated the power in this approach to facilitate prediction and/or validation of functional genomics hypotheses^17^.

We designed this study to i) determine the chromatin landscape changes that support *in vitro* hematopoiesis, and ii) determine whether *in vitro*-derived hematopoietic cells could efficiently predict complex disease loci. We used paired single-cell RNA sequencing (scRNA-seq) and single-cell assay for transposase-accessible chromatin sequencing (scATAC-seq) to profile endothelial and hematopoietic populations derived from iPSCs. We then integrated these single cell analyses with Hi-C-derived chromatin interaction data from iPSC-derived stromal/endothelial and hematopoietic cells. This multimodal platform allowed us to define dynamic transcriptional programs, transcription factor activities, and chromatin regulatory landscapes that collectively facilitate the progression from endothelial to hematopoietic cell identity. Additionally, these data uncovered evolving cell-cell interaction networks that support *in vitro* HPC emergence. In many cases, our findings mirror functional genomic regulation that occurs during *in vivo* hematopoiesis. Finally, we link *in vitro* blood cell chromatin regulation to human genetic variation with anthropometric, cardiovascular, cardiometabolic, autoimmune, neuropsychiatric, and quantitative human blood traits. These findings provide a regulatory map of chromatin regulatory elements that facilitate *in vitro* hematopoiesis, and enable variant-to-gene mapping to highlight the role of endothelial and hematopoietic cells on human phenotypes.

## Results

### Multimodal analysis of in vitro hematopoiesis

We cultured induced pluripotent stem cells (iPSCs) to model definitive hematopoiesis (**Fig. 1A**, **Supplemental Fig. S1A-B**)^19,20^. This model has been used to recapitulate hematopoiesis and a similar platform recently produced engraftable hematopoietic stem cells^13^. In our cultures, we isolated and aggregated CD34^+^ endothelial cells on day 9, and subsequently cultured in hematopoietic cytokines. Between day 9+7 and day 9+8, hematopoietic progenitor cells (HPCs) emerge from the endothelial layer directly from specialized hemogenic endothelial cells (HECs). We confirmed that *in vitro*-derived HPCs were competent to form erythroid, white cell, and megakaryocyte lineages by colony formation assays (**Supplemental Fig. 1C-D**). Thus, the endothelial-to-hematopoietic transition (EHT) occurs between day 9+7 and day 9+8.

**Figure 1.**
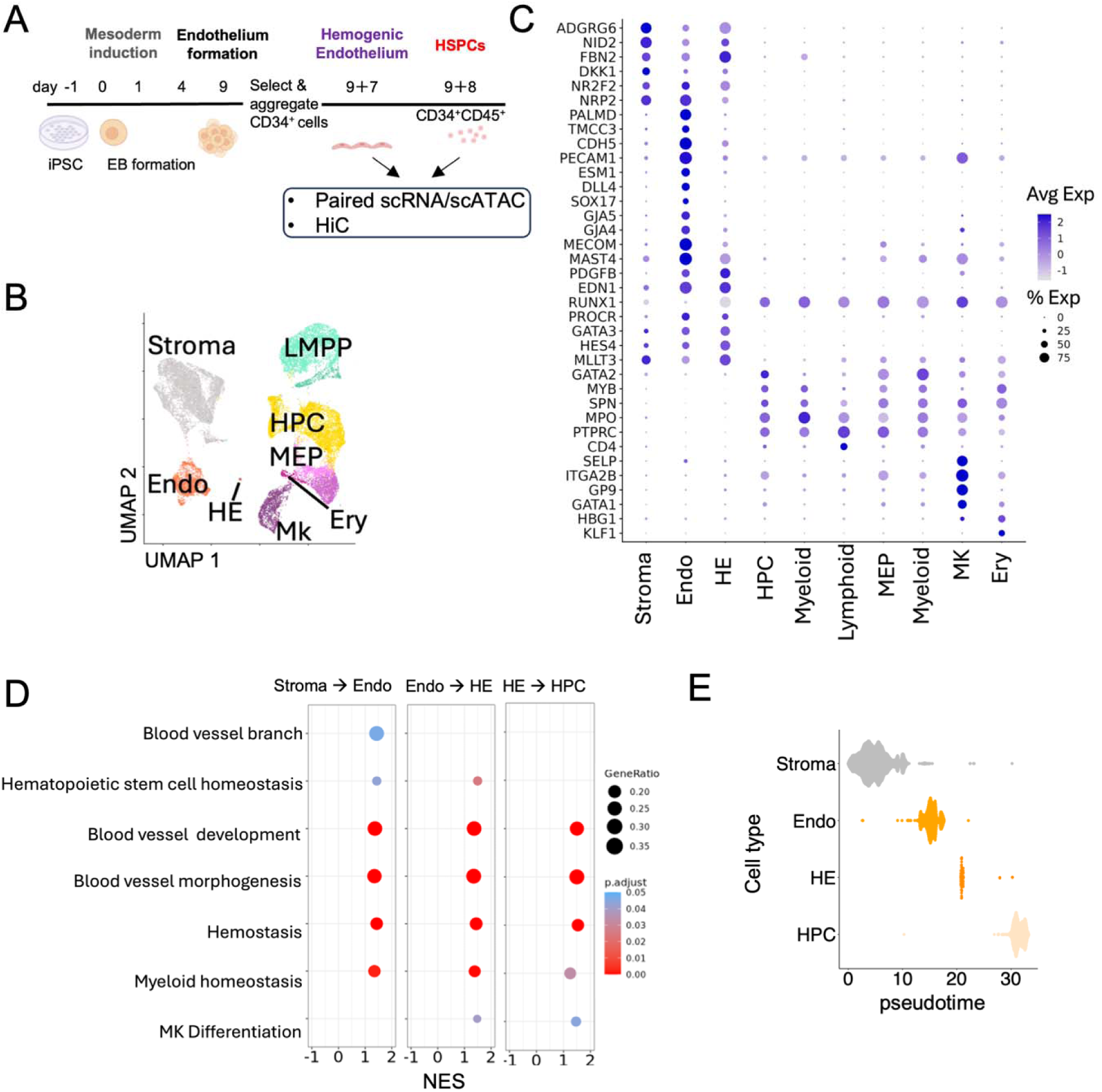
Stage-specific *in vitro* modeling recapitulates hematopoietic development. A. Schematic of the *in vitro* model and modes of analysis used in this study. B. UMAP with cell annotations and feature plots depicting key genes expressed in endothelial and hematopoietic cells. C. Dot plot identifying key gene markers supporting cell annotations. D. Pathway analysis supporting progression toward hematopoietic phenotypes in stromal, endothelial, and hematopoietic stem and progenitor cells (HSPCs). E. Pseudotime trajectory analysis supports progressive development of stromal cells and endothelia into HSPCs via endothelial-to-hematopoietic transition.

We then profiled adherent cells on day 9+7 and CD34^+^CD45^+^ non-adherent cells on day 9+8 to capture this developmental transition (**Fig. 1A**). We used a multimodal single cell approach to define adherent and non-adherent cell types at this stage, using paired single cell RNAseq/ATACseq to profile gene expression and chromatin accessibility in these cells. We further using Hi-C to ascertain chromatin interactions across adherent cells and non-adherent cells in bulk. We identified stromal, endothelial, HE, HPC, and lineage-committed hematopoietic cells produced from cultured iPSCs based on gene expression profiles (**Fig. 1B-C** and **Supplemental Fig. S2**). Given the directed differentiation protocol, in which we cultured endothelial cells in hematopoietic cytokines, endothelial cells expressed some hematopoietic genes (e.g., *MECOM*, **Fig. 1C**). However, HE cells expressed higher levels of key hematopoietic commitment genes, such as *RUNX1* and *MLLT3* (**Fig. 1C**). The relative paucity of annotated HE cells, and developmental context for our analysis during EHT, suggests that the HE cell population may be relatively late-stage cells actively undergoing EHT morphogenesis. Pathway analysis indicated an induction of hematopoietic stem cell homeostasis, platelet function, and myeloid homeostasis genes in endothelial cells, in addition to the expected increases in blood vessel morphogenesis and development associated with endothelial functions (**Fig. 1D**). These supported the idea that hematopoietic differentiation is occurring within the cultured endothelial population vs stromal cells. Compared to endothelial cells, HE cells and HPCs showed further progressive induction of hematopoietic commitment genes and processes (**Fig. 1D**).

We wondered if the CD34^low^ “stromal” population represented relatively immature endothelial cell precursors, or if these cells were instead a mature population that emerged in culture. Vascular endothelial and stromal cell types are necessary for hematopoiesis *in vivo*, providing key growth factors and ligands through direct and secreted signals in the hematopoietic niche^21^. Using trajectory analysis, we found that some stromal cells undergo transition to endothelial cells, which induce a hematopoietic transcriptional program (**Fig. 1E**). However, we could not exclude the possibility that some stromal cells captured in our analysis represent distinct, mature cell types with roles similar to vascular niche cells *in vivo*. This finding suggests that the definitive iPSC culture system may produce stromal and endothelial cells necessary to reconstruct a complex hematopoietic niche, in addition to *bona fide* HE cells and HPCs.

Trajectory analysis also indicated distinct developmental trajectories for major hematopoietic lineages, including lymphoid-, myeloid-, megakaryocytic-, and erythroid-based progenitor cells with lineage-specific gene expression markers (**Fig. 1B, 1E**). Note that these lineage-based CD34^+^CD45^+^ progenitor cells do not represent mature cell types, although they do retain an ability to produce more mature blood cell types in hematopoietic functional assays (**Supplemental Fig. S1**).

### Transcriptional regulatory activities dictate hematopoietic lineage commitment in vitro

We next assessed chromatin accessibility and transcriptional activities during *in vitro* EHT. We quantified the number of accessible chromatin peaks at single cell resolution (scATACseq), and ascertained cis-regulatory regions (cREs) involving 3-dimensional chromatin interactions by HiC^17^ (**Supplemental Figs. S2-S3**). We assigned cREs based on Hi-C contacts within accessible chromatin regions (scATACseq peaks), permitted cell type-specific interpretation. We found that chromatin accessibility and cRE activities were high in stromal (n=206,905 ATAC peaks, 24,399 cREs), Endo (n=122,723 ATAC peaks, 14,511 cREs) and HPC populations (n=136714 ATAC peaks, 22325), and that chromatin accessibility generally decreased during initial lineage commitment (**Fig. 2A**). For example, Mk-biased progenitors (n=85,993 ATAC peaks, 15,249 cREs) and erythroid-biased progenitors (n=45,158 ATAC peaks, 7,902 cREs) contained the fewest open chromatin regions of the cells analyzed (**Fig. 2A**). This is consistent with major chromatin reorganization occurrence at the onset of hematopoietic lineage commitment^22,23^. The relative paucity of open chromatin regions and cREs in HE cells may be due to developmental changes or alternatively could be related to having captured relatively few HE cells.

**Figure 2.**
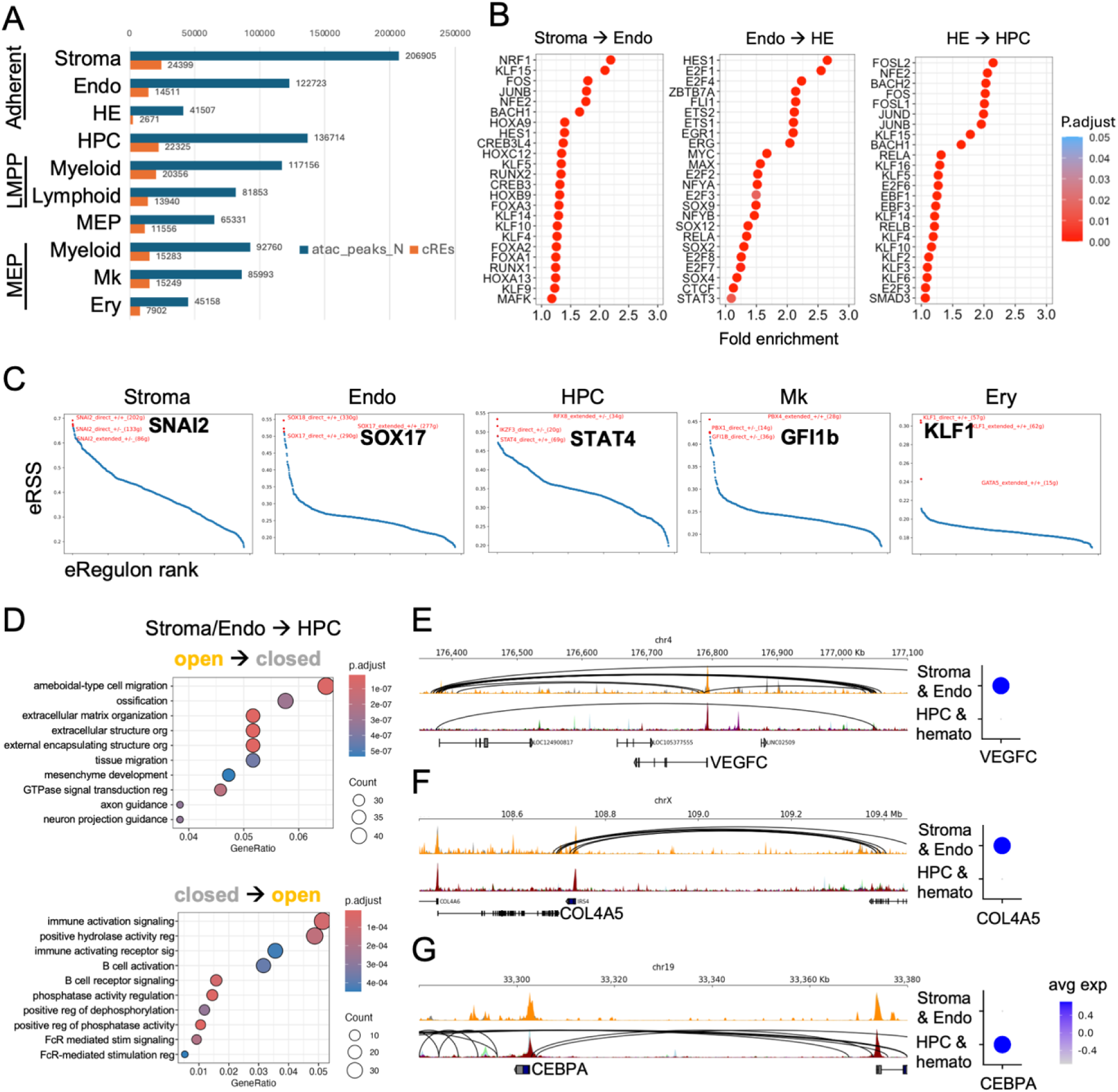
Single cell multiomics defines transcription factors and genomic regulatory elements underlying *in vitro* hematopoiesis. A. Number of accessible chromatin regions (ATACseq peaks) and active cis-regulatory elements (cREs) as defined by Hi-C in the indicated cell types. Dashed line separates adherent and non-adherent cell types. B. Differentially accessible TF binding sites as defined by comparing the indicated cell types. Fold enrichment reflects increased accessibility in Endo, HE, or HPCs as indicated in the panels. C. Active expression regulatory networks (eRegulons) driven by the indicated transcription factors in each cell type. D. Pathway analysis results for genes that open or close during the transition from adherent to non-adherent cells. Dot size reflects the overall gene count and color depicts statistical significance. E. The VEGFC locus is active and participates in 3D chromatin interactions in adherent cells, but these interactions are eliminated during EHT. F. Extracellular matrix genes and related regulatory molecules are are reduced during EHT. The COL4A5 gene promoter contains active chromatin regulatory loops in adherent cells, which are eliminated in non-adherent hematopoietic cell types. G. The CEBPA gene promoter is accessible but quiescent in adherent cell types and becomes actively engaged in chromatin loops in non-adherent hematopoietic cells after EHT.

We then performed transcription factor (TF) motif enrichment analysis across *in vitro* cell types to identify transcriptional regulators for *in vitro* hematopoiesis (**Fig. 2B**). Analysis of common TFs with increased accessibility in endothelial (Endo) vs stromal cells included binding sites for TFs that regulate hematopoietic commitment, such as RUNX1, HOXA9, and NFE2 (n=420 statistically significant changes). These findings were consistent with the notion that the *in vitro* Endo population are committed to a hematopoietic fate. Differences between Endo and hematopoietic endothelial (HE) cells showed an enrichment in accessible sites for TFs known to mediate EHT, including HES1 (Notch pathway) and MYC (n=238 significant changes). Finally, comparison of HE and hematopoietic progenitor cell (HPC, n=350) chromatin accessibility revealed increased accessibility ta bindings sites for TFs known to be associated with hematopoietic stem cell (HSC) maintenance, differentiation, and lineage commitment, including NFE2, JUNB/D, and EBF1/3. In sum, changes in chromatin accessibility reflected sequential differentiation and developmental transitions during EHT and hematopoietic fate commitment.

We next sought to connect TF activities to gene expression regulation during *in vitro* differentiation. We assessed regulon specificity scores (RSSs) in stromal, endothelial, HPC, HE, and hematopoietic cell types^24^ (**Supplemental Fig. S2B-C**). Cell type-specific gene expression patterns were consistent with key TF activities. In stromal and endothelial cells, the most enriched RSSs were linked to SNAI2 and SOX17 activities, respectively (**Fig. 2C**). SNAI2 and SOX17 play essential roles in epithelial to mesenchymal transitions (EMT)^25^ and EHT^26^, respectively. Active SNAI2 and SOX17 transcriptional programs align with the developmental changes ongoing in our iPSC system at the time of cell collection (**Fig. 1A**). Active SNAI2 signaling in stromal cells also supported the idea that some CD34^low^ cells within the stromal population are undergoing developmental transition into endothelial cells, consistent with trajectory analysis (**Fig. 1E**). Other regulon activities reflected known and novel factors that regulate hematopoietic identify and development. For example, the top RSS in HPCs was STAT4. STAT4 activity regulates HPC homeostasis *in vivo* ^27^. Furthermore, GFI1b and KLF1 are crucial for megakaryocytic and erythroid fate determination, respectively, and were enriched in Mk and Ery committed progenitors *in vitro* (**Fig. 2C**) ^22,23^.

Genome-wide, there were similar numbers of loci with dynamic accessibility across EHT (**Supplemental Fig. S3A-D**). cREs that disappeared during EHT (n=1652 loci AtoB) as interaction sites that appeared during EHT (n=1740 loci BtoA). Gene ontology analysis of the sites that closed (AtoB) showed enrichment of genes related to cell migration and extracellular matrix organization (**Fig. 2D** and **Supplemental Fig. S3E**). These processes are integral to endothelial cell identity and EHT^28^. In contrast, loci and genes related to immune activation and signaling became accessible and active in HPCs after EHT (**Fig. 2D** and **Supplemental Fig. S3E**). A ‘sterile’ inflammatory response is required for EHT and initial HPC production^29^. The widespread chromatin reorganization identified in this experiment is indicative of the dramatic changes in morphology and cell state that occurs during EHT, but argues against global chromatin shutdown or activation during this process.

Dynamically regulated cREs during EHT indicated developmental changes. When we broadly compared cREs in adherent (stromal, endothelial, HE cells) vs non-adherent progenitors, we identified key loci related to cell identity. For example, vascular endothelial and stromal interaction factors were shut down during EHT. Adherent cells showed cREs in the promoter regions for *Vascular Endothelial Growth Factor C* (*VEGFC*) and *Collagen 4A5* (*COL4A5*), but chromatin interactions were lost in HPCs (**Fig. 2E-F**). These genes function in endothelial growth and stromal attachment, and support hematopoietic niche formation *in vivo*^21^.

Although *in vitro* endothelial cells showed some induction of the hematopoietic transcriptional program (**Fig. 1**), we noted new chromatin interactions at key hematopoietic loci like *CCAAT enhancer-binding protein alpha* (*CEBPA*) in HPCs (**Fig. 2G**). *CEBPA* is expressed in nascent HPCs and is essential for their generation and maintenance^30,31^. In sum, these findings add insight into chromatin regulatory events that accompany EHT, including biological processes that enable the dramatic morphologic and developmental changes necessary for nascent HPC formation.

### Intercellular communication promotes hematopoietic differentiation in vitro

We hypothesized that stromal and vascular endothelial cells might support HE and HPC development *in vitro*, as they do *in vivo*^21^. We ascertained active cell-cell signaling among *in vitro* cell populations using CellChat^32^. We broadly identified ligand-receptor pairs known to promote EHT *in vivo*, including ANGPTL, PROS, and PTN (**Fig. 3A**). This suggests that similar signaling mechanisms exist *in vitro* as they do *in vivo*, and indicates that *in vitro* adherent cells may promote EHT and HPC production. In fact, adherent cells secreted and/or responded to a number of signals that regulate EHT (**Fig. 3B-G**). TGFb signaling is a key factor that promotes EHT^33,34^. We identified active TGFb signaling from adherent cells that was specifically received by TGFβ receptors on HE cells (**Fig. 3B, 3E**). Bone morphogenic protein (BMP) signaling is critical to endothelial specification and EHT *in vitro*^35,36^. The BMP ligands GDF6 and GDF7 secreted from endothelial cells were received primarily by HE cells *in vitro* (**Fig. 3C, 3F**). We also noted metabolic signals emerging from adherent populations that may support EHT and/or hematopoietic cells. IGF and IGFBP signaling were active *in vitro* (**Fig. 3A**), and IGFBP3 secreted from stromal cells was received specifically by HE cells (**Fig. 3D, 3G**). This ligand-receptor interaction may regulate cell survival in the setting of IGF1 and other growth factors in culture^37^.

**Figure 3.**
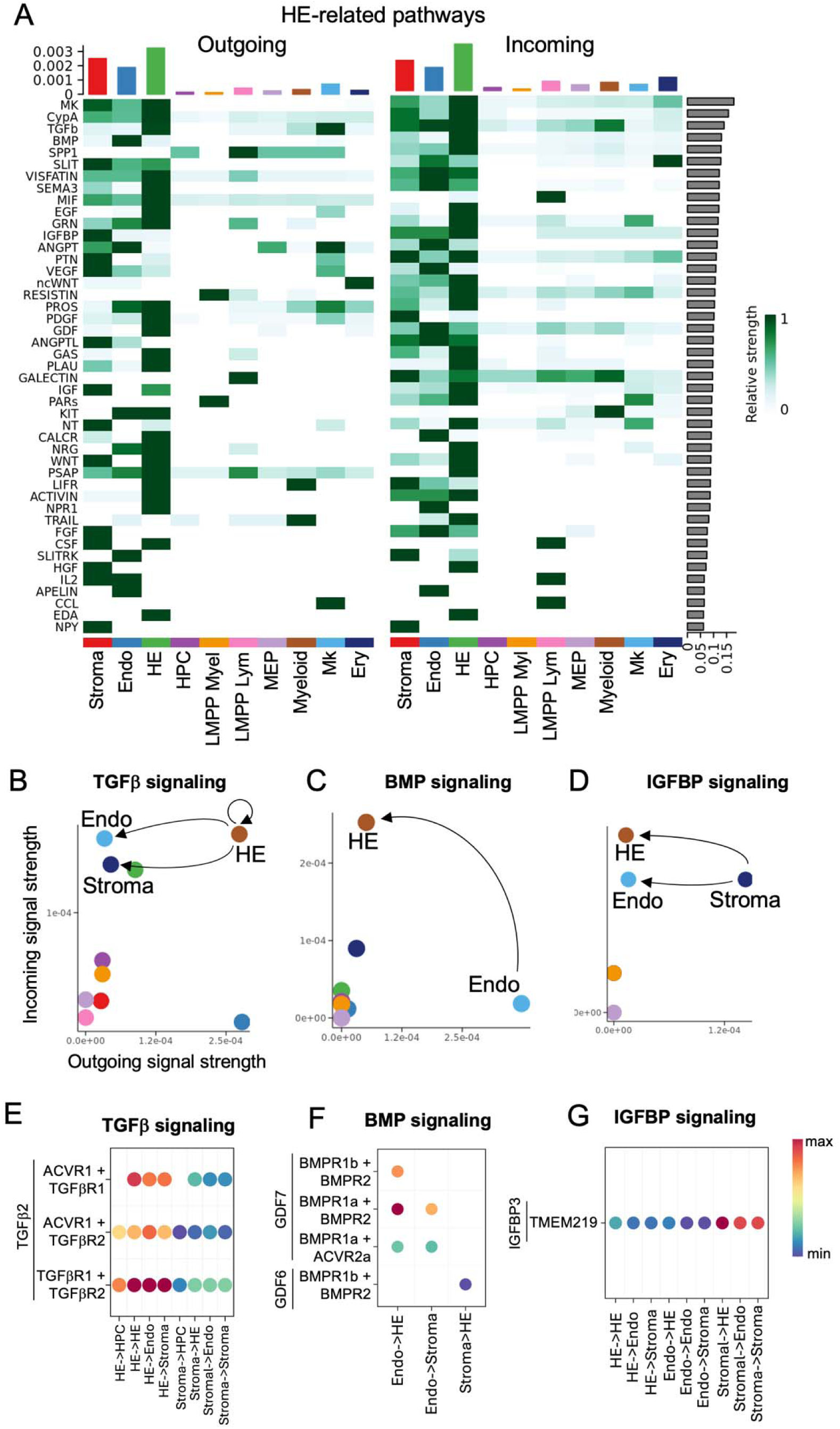
Cell-cell interaction networks impact *in vitro* EHT. A. Top ligand-receptor networks active in HE cells. Axes represent numerical confidence estimates for signaling activities, relative to other cell types. B. TGFβ signaling activities during *in vitro* hematopoietic culture. Dots indicate individual cell types. Ligands secreted by HE and Mk progenitor cells are received by Stroma, Endothelial, and HE cells. C. BMP signaling activities during *in vitro* hematopoietic culture. Dots indicate individual cell types. Ligands secreted by Endothelial cells are received by HE cells. D. IGFBP signaling activities during *in vitro* hematopoietic culture. Dots indicate individual cell types. Ligands secreted by Stroma are received by Endothelial and HE cells. E-G. Identities for most active ligand-receptor interactions for the indicated signaling pathways. Colors reflect algorithm confidence scores for interactions.

We also noted active inflammatory and immunoregulatory signals emanating from hematopoietic progeny cells at day 9+8, including progenitors biased toward immune lineage fates. TRAIL, SPP1/Osteopontin, and PAR ligands secreted by circulating immune cells *in vivo* are normally received by endothelial cells and promote HE formation via sterile inflammation^21,38^. We noted analogous signals from nascent lymphoid and myeloid-biased progenitors, which were also received by surface receptors on *in vitro*-derived HE cells (**Fig. 3A**). These sterile inflammatory signals promote formation of definitive engraftable HPCs *in vivo*, and may play similar roles *in vitro*. In sum, our paired single cell RNA/ATAC data confirmed that the *in vitro* system recapitulated known signals that direct developmental hematopoiesis with a hematopoietic niche that forms spontaneously during *in vitro* culture.

### In vitro hematopoiesis as a model for understanding genetic regulation of complex human traits

Finally, we wanted to leverage our multimodal single cell analysis of *in vitro*-derived blood cells to help define genetic determinants of complex disease. We used partitioned linkage disequilibrium score regression (LDSR) on open chromatin regions for each cell type to ascertain cell-specific GWAS signal enrichment^17^. *In vitro* cREs for *in vitro* stromal, endothelial, and hematopoietic cell types effectively marked loci related to complex human trait variation (**Fig. 4A**).

**Figure 4.**
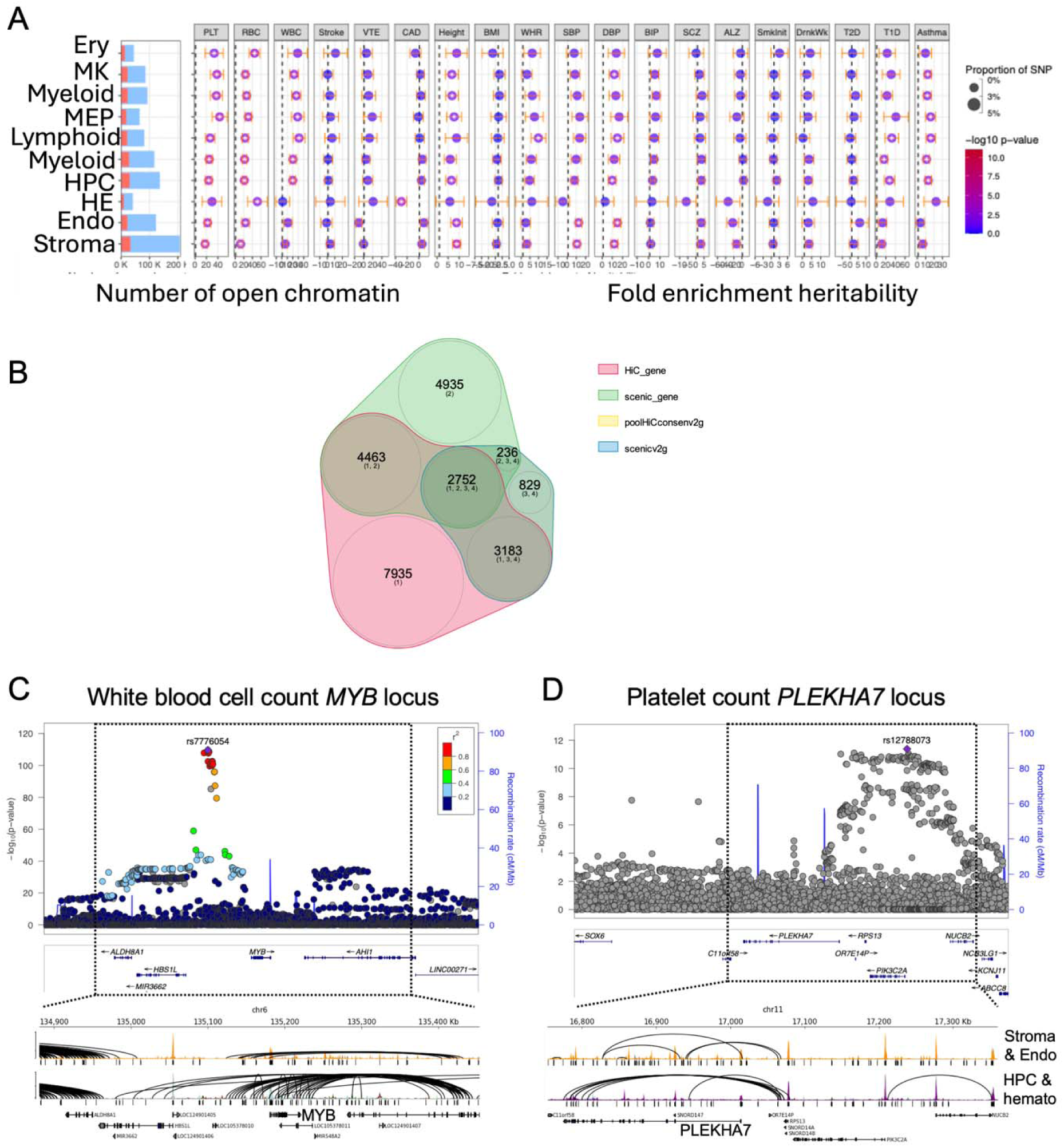
Active regulatory loci ascertained using *in vitro*-derived hematopoietic cells predict complex human disease traits. A. *In vitro* cell types predict human traits. Open chromatin regions (blue) and active cis-regulatory elements (cREs, orange) are shown. Forest plots depict heritability enrichment by LDSC analysis for each cell type on the indicated disease or trait. Open dots reflect significant associations (p<0.05). Vertical dashed lines in each plot indicate line of no enrichment (heritability = 1). Dot sizes indicate the proportion of contributed heritability to each trait. B. Venn diagram depicting overlaps in variant-to-gene predictions from four separate models. C. Exemplary *MYB* gene locus highlights cell-specific chromatin interactions at a GWAS locus for white blood cell count. Locus zoom plot shows genome-wide significant SNPs in the intergenic region between *MYB* and *HBS1L*. Tracks below show accessible chromatin peaks and chromatin interactions for adherent (top) and non-adherent (bottom) cell types. Peaks are shown below tracks, identifying additional chromatin peaks and chromatin interactions in HPCs and white blood cell-based progenitors (Lymphoid and Myeloid) in the intergenic region overlapping GWAS signal. D. Exemplary *PLEKHA7* gene locus highlighted cell-specific chromatin interactions at a GWAS locus for platelet count. Locus zoom plot shows genome-wide significant SNPs near the *PLEKHA7* gene, which also overlap the *PIK3CA* gene locus. Tracks below show accessible chromatin peaks and chromatin interactions for adherent (top) and non-adherent (bottom) cell types. Peaks are shown below tracks, identifying additional chromatin peaks and chromatin interactions in HPCs and megakaryocyte-biased progenitors. Note separate loci that participate in chromatin loops upstream of PLEKHA7 and others that contact the PIK3CA promoter, suggesting separable GWAS signals.

We cross-referenced GWAS loci from anthropometric, cardiovascular, cardiometabolic, neuropsychiatric, behavioral, autoimmune, and quantitative blood traits to determine genetic relationships between accessible chromatin regions from *in vitro* blood cells with these complex human traits ^39,42,49,58,60,62,64,66,68–73^. As expected, these experiments revealed enrichment for loci related to quantitative platelet, erythroid, and white blood cell traits among accessible regions in our *in vitro*-derived hematopoietic cells (**Fig. 4A**). Notably, there was dynamic enrichment for some traits, including WBC loci, demonstrating specificity in these findings. Stromal and endothelial cell types were *not* enriched for WBC loci, whereas hematopoietic cell types showed significant WBC locus enrichment. We noted a similar pattern for (auto)immune traits, including asthma and type 1 diabetes (T1D), which only showed enrichment linked to hematopoietic cell types (**Fig. 4A**). In contrast, blood pressure traits (e.g., systolic blood pressure, SBP) showed significant association with stromal and endothelial cell types and not hematopoietic traits. These findings underscore the utility for *in vitro*-derived cells to model tissue-specific *in vivo* biology.

### In vitro hematopoietic cells identify dynamic chromatin regulation at loci implicated in blood trait variation and complex disease inheritance

The significant enrichment of GWAS loci among *in vitro*-derived cells motivated variant-to-gene mapping for select traits. We employed four scenarios to identify genes related to cREs, including Hi-C, SCENIC, poolHiCconsenv2g, and scenicv2g (**Fig. 2B**). A total of 2,752 locus-gene pairs were identified by *all* four methods. Targeted evaluation of these loci revealed dynamic genomic regulation across *in vitro*-derived cell types and highlighted genes and mechanisms to explain quantitative blood traits and blood cell-related disease pathology.

A canonical intergenic region harbored between the *HBS1L* and *MYB* genes has been associated with blood cell traits across lineages, including white blood cell count (**Fig. 4C**). This locus showed accessible chromatin regions in lymphoid and myeloid cells derived in our assays, with chromatin interactions at the *MYB* promoter that are well positioned to regulate gene expression (**Fig. 4C**). We noted increased chromatin accessibility and looping in HPC and hematopoietic cells, as compared to adherent stromal and endothelial cells, in keeping with the dynamic regulation of this locus and others that regulate white blood cell count (**Fig. 4A**).

We also resolved complex platelet trait loci through our analysis. The *PLEKHA7/PIK3CA* gene locus has been implicated in quantitative platelet count variation, among other platelet traits ^39,40^ (**Fig. 4D**). Most significant SNPs are upstream of the *PLEKHA7* gene locus, overlapping the *PIK3CA* gene body. Within this locus, we identified looping events between GWAS SNPs and the *PLEKHA7* gene promoter, as well as the *PLEKHA7* gene body (**Fig. 4D**). The open chromatin regions and looping events also changed between stromal/endothelial cells and HPCs, including a novel chromatin interaction between GWAS SNPs and the *PIK3CA* gene promoter in HPCs (**Fig. 4D**). This finding implicates regulation of both *PLEKHA7* and *PIK3CA* genes in genetic activities at this locus, which are predicted to occur within hematopoietic cells generated in culture.

We further probed variant-to-gene predictions related to human disease states. Our LDSC findings showed that *in vitro*-derived hematopoietic cells, but not stromal or endothelial cells, were enriched for open chromatin in autoimmune disease trait loci (**Fig. 4A**). For example, the *IL1RL1* gene locus has been implicated in Asthma ^41^ (**Fig. 5A**). Examination of this locus within our data revealed extensive chromatin regulation at the *IL1RL1* gene promoter and intronic regions (**Fig. 5A**). Most of the chromatin contacts at this site emerged in hematopoietic cells, particularly lymphoid- and myeloid-biased progenitors, as opposed to stromal and endothelial cells (**Fig. 5A**). These findings directly link asthma predisposition SNPs with *IL1RL1* gene regulation, in addition to nearby *IL18R1* and *IL1R1*.

**Figure 5.**
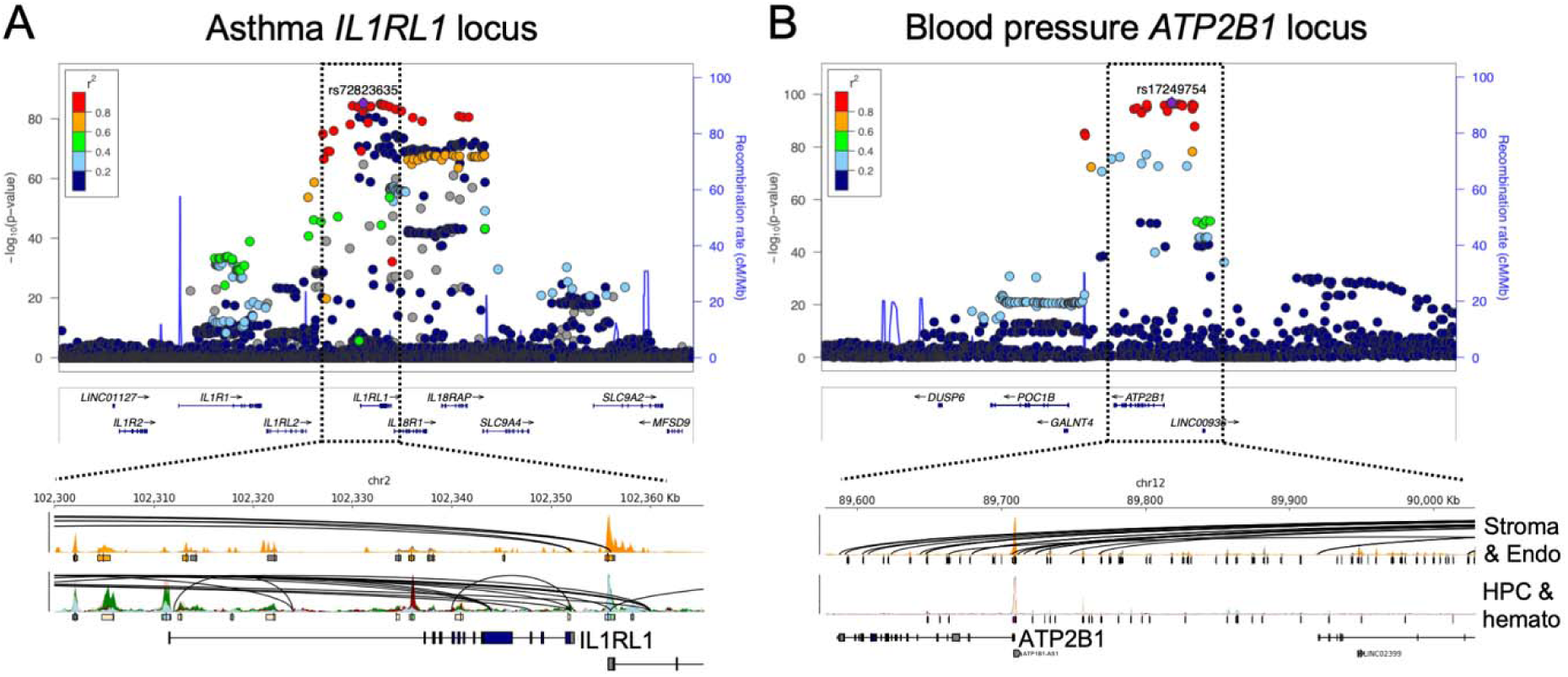
*In vitro*-derived hematopoietic cells predict asthma and blood pressure regulatory loci. A. *In vitro* cells implicate regulatory SNPs for *IL1RL1* in asthma predisposition. LocusZoom plot shows significant loci mediating asthma risk at the *IL1RL1* locus. Boxed area is highlighted at bottom, showing chromatin looping interactions at open chromatin regions within the *IL1RL1* promoter and gene body that are specific to hematopoietic cell types, including myeloid and lymphoid-biased progenitors. B. *In vitro* cells implicate regulatory SNPs for *ATP2B1* in blood pressure modulation. LocusZoom plot shows significant loci mediating systolic blood pressure at the *ATP2B1* locus. Boxed area is highlighted at bottom, showing chromatin looping interactions at open chromatin regions within the *ATP2B1* promoter and gene body that are specific to adherent stromal and endothelial cell types. These interactions are abrogated in hematopoietic cells.

Finally, we interrogated the finding that stromal and endothelial cells were enriched for open chromatin at sites related to blood pressure trait variation, whereas HPCs and hematopoietic cells did not show enrichment for these traits (**Fig. 4A**). Among variant-to-gene predictions related to blood pressure traits were SNPs related to *ATP2B1*, which has previously been implicated in blood pressure variation (**Fig. 5B**). Stromal and endothelial cells were indeed enriched for open chromatin at the *ATP2B1* promoter, gene body and upstream regions (**Fig. 5B**). Perturbation in these long-range chromatin interactions would impact *ATP2B1* expression in endothelial cells, directly implicating this gene as relevant for the blood pressure variation noted in prior GWAS ^42^.

## Discussion

The objectives of this study were to determine chromatin landscape changes that accompany the endothelial-to-hematopoietic transition in a human cell model, to define genetic regulatory elements and signaling that are active during *in vitro* EHT, and to test if an *in vitro* hematopoietic system can effectively model genetic variation known to impact human blood traits and complex disease susceptibility. Using a multimodal single cell approach, we ascertained relevant gene expression changes, transcription factor activities, and pathway changes during *in vitro* EHT. These changes effectively mirrored *in vivo* developmental hematopoiesis from endothelial specification through hematopoietic lineage commitment, providing new depth in understanding *in vitro* hematopoiesis that could help support production of translationally useful cell types.

Our study elucidated cell signaling modalities that support *in vitro* HPC formation, some of which mirrored signaling modalities that also support *in vivo* hematopoiesis. In addition to established TGFβ and BMP signals, our findings implicate metabolic and apoptotic signaling mechanisms in the development of HPCs *in vitro* (**Fig. 3G**). Secreted IGFBP3 complexes with IGF1 to promote survival or TMEM219 to promote apoptosis^43^. Identifying this signaling interaction highlights the importance of IGF1 during EHT and HPC formation. Although we and others view stromal interactions as generally supportive to HPC emergence^21^, these findings highlight a potentially counterproductive stromal effect on *in vitro* hematopoiesis, since stromal cells provide the majority of IGFBP3 in our culture system. These developmental insights into the *in vitro* system have implications for moving *in vitro* blood cell production toward clinical translation. The intercellular signaling modalities elucidated in this study can be used to modify or augment the *in vitro* hematopoietic platform to produce HPCs and hematopoietic cells to support a range of basic and translational avenues.

A major goal of this study was to ascertain dynamics in the chromatin landscape during iPSC-derived blood cell production. In addition to developmental hematopoiesis insights, defining open chromatin regions and cis-regulatory elements insights help predict loci related to human blood trait variation and a range of heritable phenotypes (**Fig. 4**). Our results indicate that the chromatin environment in stem cell-derived HPCs specifies variants and genes relevant for quantitative blood trait variation, autoimmune disease pathology, and in some cases anthropometric traits (e.g., height and blood pressure, **Fig. 4A**). These interactions were also relatively trait-specific, with non-significant associations noted between our *in vitro*-derived cells and several cardiovascular, neuropsychiatric, and behavioral traits (**Fig. 4A**). The dynamic changes in chromatin state between adherent and non-adherent *in vitro*-derived cells was remarkable and permitted us to evaluate dynamic chromatin regulation in our data set. For example, the *MYB*, *PLEKHA7/PIK3CA*, and *ATP2B1* loci all represent regions with significant changes in accessibility and chromatin interactions (**Figs. 4-5**).

Strong *in vivo*-*in vitro* correlates at the level of single cell chromatin accessibility suggest that human genetic insights might be used to augment *in vitro* cell production. There remains strong interest in producing blood cell-based therapeutics for clinical testing^44^, transfusion medicine^28,45^, and other cell therapeutics^46^. One major barrier to clinical translation is inefficient EHT and blood cell production. We demonstrate that the definitive *in vitro* system recapitulates *in vivo* developmental biology with regard to EHT and hematopoietic lineage commitment (**Fig. 1-3**). Given links to human genetic variation, including quantitative blood trait variation, it is plausible that human GWAS loci can inform locus and/or gene modifications that boost blood cell production *in vitro* as they do *in vivo*^10,11^.

Additionally, there is considerable interest in the directed differentiation of cells to support production of disease-modifying blood cell-based therapeutics (e.g., regulatory immune cells for asthma). Loci revealed through our study provide a wealth of potential loci and mechanisms for this purpose, including the important contributions of stromal cells in the hematopoietic niche that may direct hematopoietic specification (**Fig. 3**)^21^.

Our findings also indicate that *in vitro* models can help define genomic and biological regulatory activities. For example, we identified enrichment of autoimmune disease-related loci to be accessible in hematopoietic cell types, and blood pressure loci conversely predicted by stromal and endothelial cell types. It will be interesting to interrogate chromatin dynamics in rare or hard-to-isolate human blood cell types to help explain complex genetic disease susceptibility. For example, brain microglia or other tissue-resident macrophages cannot be easily isolated in adequate quantities from primary tissue but can be produced in culture. Neurodevelopmental and psychiatric phenotypes like autism may thus be better understood through multimodal profiling of these *in vitro*-derived blood cells, using the approach taken herein.

Collectively, our results provide compelling evidence that iPSC-based models of human hematopoiesis capture key aspects of the regulatory architecture governing blood cell development and trait variation. These models represent a powerful and scalable platform to dissect gene regulatory mechanisms, interpret GWAS findings in appropriate cellular contexts, and uncover previously unrecognized pathways relevant to blood disorders and complex human disease states. Integrating multimodal single cell profiles with 3D genome architecture in the *in vitro* context can guide precision medicine development and accelerate the translation of genetic discoveries into clinical applications.

## Acknowledgements

The authors thank the Children’s Hospital of Philadelphia Center for Applied Genomics for assistance with single cell library preparation and sequencing.

## Methods

### Single cell sample preparation

We performed definitive hematopoietic differentiation and nuclear isolation as described^19,20^, using pooled adherent cells on day 9 + 7 and FACS-sorted CD34^+^CD45^+^ hematopoietic cells on day 9+8. Library preparation followed Chromium Next GEM Single Cell Multiome ATAC + Gene Expression platform protocols (10x Genomics). Libraries were sequenced on an Illumina NovaSeqTM 6000 and processed using standard packages and workflows^47,48,50^ (see **Supplemental Methods**). Antibodies used for these studies are listed in **Supplemental Table S10**.

### Gene set enrichment and gene ontology analysis

Gene expression was defined for each cluster using the FindMarkers function (Seurat v5) and enrichment was assessed using the Molecular Signature Database (MSigDB) ‘Hallmark’, ‘C2’, ‘C3’, and ‘C5’ gene sets (fgsea)^51^. Pairwise comparisons for indicated cell types are presented as Normalized Enrichment Scores (NES). We used the clusterProfiler R package to perform gene ontology analysis on genes that were opened or closed during hematopoiesis^52^.

### Trajectory analysis

Slingshot and Monocle3 were used to assess trajectory analysis using default parameters^53,54^ (**Supplemental Methods**).

### Cell-cell interaction analysis

We used CellChat (v2.1.1) with default parameters to assess cell-cell communication^32^ (see Supplemental Methods).

### Open chromatin region and cRE identification

Integration of paired scRNA/ATACseq data and Hi-C were completed essentially as described to identify cell type-specific open chromatin regions and chromatin looping interactions, from which we defined cis-regulatory elements^17^. We assigned eRegulon specificity scores using SCENIC+^24^.

### Motif Identification and Analysis

Motif identification and enrichment analysis were performed on differentially accessible peaks between Endo, HE, HPC, and stromal cells using the Seurat and JASPAR2020 R packages. Motif position frequency matrices (PFMs) from the JASPAR CORE collection for vertebrates were retrieved using the getMatrixSet function, and these motifs were incorporated into the Seurat object with the human genome reference (hg38) using the AddMotifs function. Differential peak accessibility between cell types was assessed using the FindMarkers function, applying a linear regression model with nCount_ATAC as a latent variable and a minimum of 5% of cells expressing each peak. Top differentially accessible peaks (p-value < 0.005) were selected for motif enrichment analysis using the FindMotifs function.

### Transcription Factor Footprint Analysis

To define transcription factor (TF) binding site enrichment on a cell type-specific level, we performed footprint analysis using the Signac R package^55^. TF motifs were retrieved from JASPAR^56^ using the getMatrixSet function and integrated into our single cell object with the AddMotifs function (Seurat v5)^57^.

### GWAS collection

Summary statistics were collected from public repositories. Details can be found in **Supplemental Table S11**^39,42,49,58,60,62,64,66,68–73^. Proceduress used to reformat and analyze GWAS, including LDSC, can be found in **Supplemental Methods.**

### Genomic locus visualization

We used pyGenomeTracks to display chromatin accessibility and chromatin looping events at key loci^59,61^. We present GWAS loci using LocusZoom^63,65^.

## Data availability

Raw sequencing data will be available on the NCBI Gene Expression Omnibus. All scripts used publicly available software packages.

