## Supplemental Figures and Table Legends for "Multimodal analysis of *in vitro* hematopoiesis reveals blood cell-specific genetic impacts on complex disease traits"

Christopher S Thom

10-052 Colket Translational Research Building

3501 Civic Center Blvd

Philadelphia, PA 19104

Struan FA Grant

1102D Abramson Research Center

3615 Civic Center Boulevard

Philadelphia, PA 19104

**Supplemental Figures and Table Legends**

**Supplemental Figures**

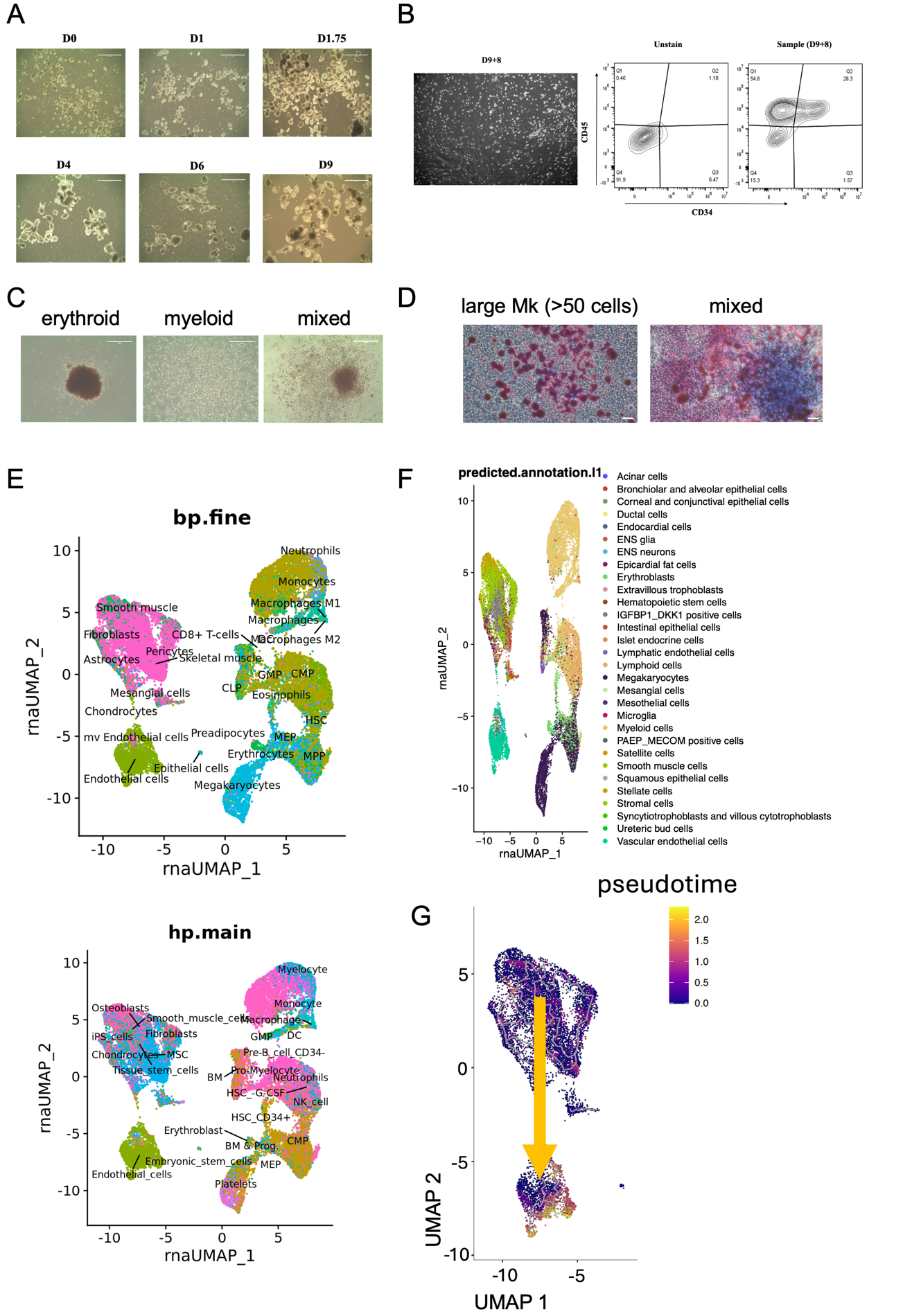

**Supplemental Figure S1.** Functional analysis and cell annotations for definitive hematopoietic cells produced *in vitro*. **Related to Figure 1**

A. Exemplary images of embryoid body development during definitive *in vitro* hematopoiesis.
B. Exemplary FACS plot for cells analyzed in this study. Day 9+8 CD34^+^CD45^+^ HPCs (shown) were sorted and analyzed along with adherent cells from Day 9+7 in this study.
C. Exemplary colonies produced by *in vitro* HSPCs, including mixed lineage colonies and large definitive erythroid colonies. Scale bar, 400 um.
D. Exemplary colonies produced by *in vitro* HSPCs, including large megakaryocyte colonies. Scale bar, 50 um.
E. SingleR cell annotations, which contributed to identification and assignment of cell types.
F. Comparison to the Azimuth data repository, which contributed to identification and assignment of cell types.
G. Pseudotime analysis of stromal and endothelial cells using Monocle3, which identifies a progressive development of some stromal cells into the endothelial cell population. These endothelial cells gain hematopoietic gene expression prior to undergoing EHT.

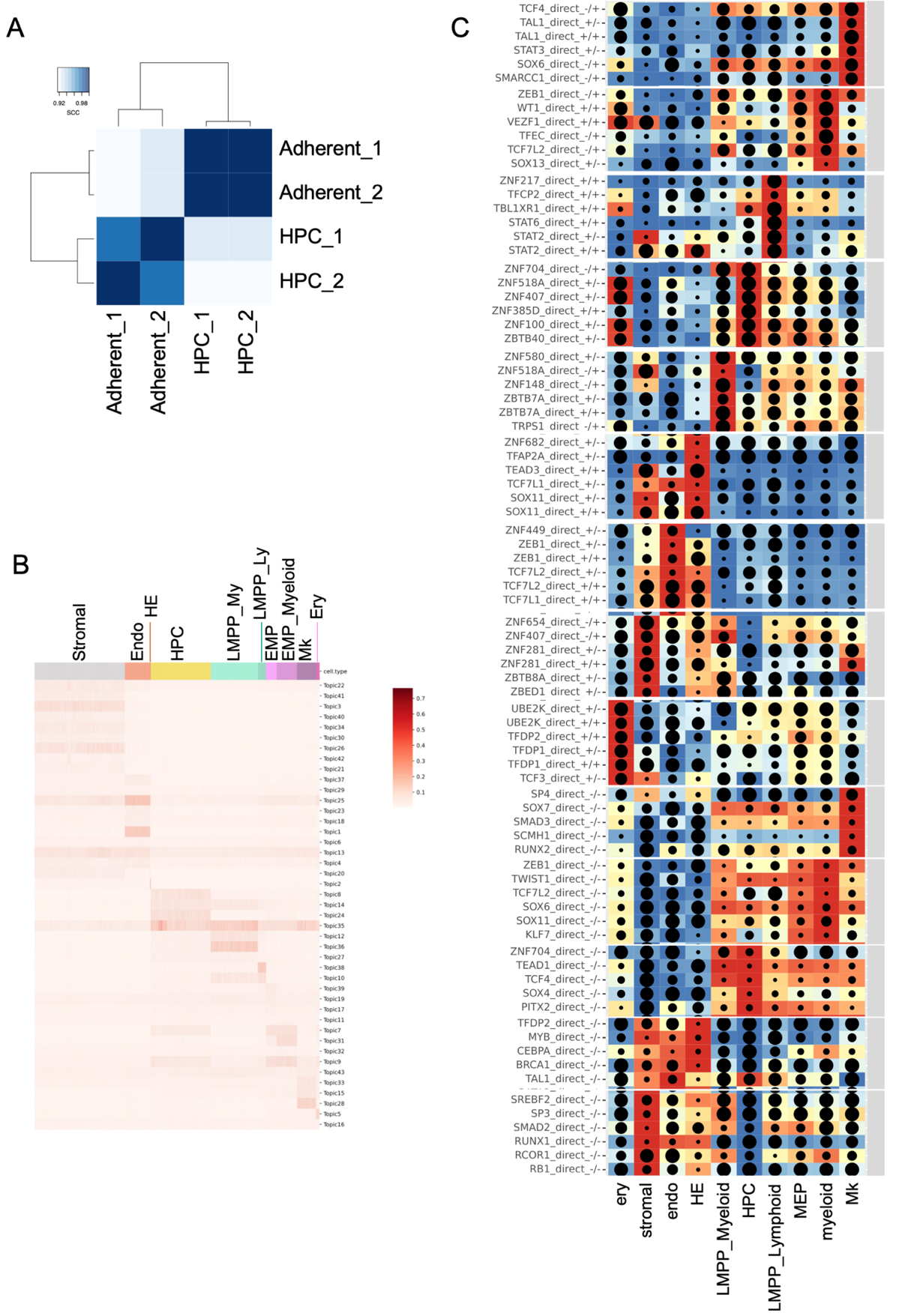

**Supp Fig. 2. Genomic correlation and transcriptional regulatory activities for *in vitro* derived cells. Related to Figure 2.**

A. Correlation plot for bulk Hi-C samples analyzed.
B. Cell type-specific ‘topic’ enrichments for *in vitro* derived cells.
C. Enriched transcription factor activities ascribed to each cell type.

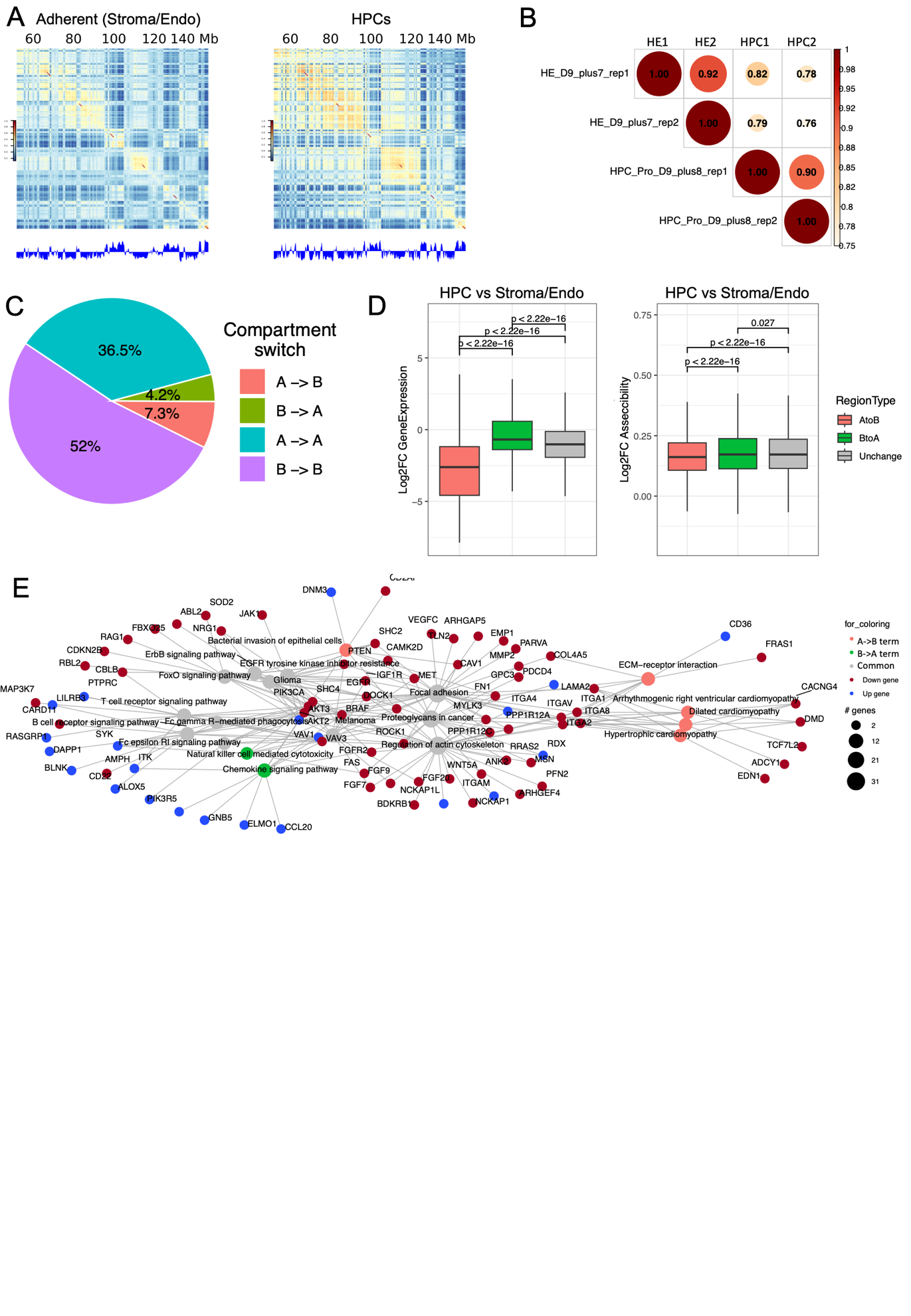

**Supp Fig. 3. Chromatin domain analysis across cell types and compartments during EHT. Related to Figure 4.**

A. We estimated A/B compartments in adherent stroma/endothelial cells and HPCs by eigenvector analysis of the genome contact matrix at 40 kb resolution after observed/expected frequency normalization. Shown are examples of Pearson correlation matrices of A/B compartment identification at 40 kb resolution on chromosome 8. The plaid pattern suggests that chromatin spatially segregates into two compartments. The first eigenvector of the correlation matrix is shown below to derive compartment type, with genomic GC content as reference.
B. Jaccard coefficient of A/B compartment assignments by pairwise comparison for each replicate sample derived from stromal/endothelial and HPC samples. A/B compartment assignment was defined by the sign of the first eigenvector for each 40 kb bin and compared between Hi-C samples. While there are differences in the A/B compartment organization between the stromal/endothelial and HPCs, there is still a reasonable degree of similarity (coefficients are well above 0.5). We noted variability across cell type replicates by associated coefficients (0.9-0.92).
C. Composition of genomic regions that changed compartment status or remained the same in adherent stromal/endothelial cells and HPCs, with 7.3 % of the compartmentalized genome opened and 4.2% becoming closed.

D-E. Distribution of fold change in (D) gene expression or (E) OCR accessibility at dynamic

(A→B or B→A) or stable (A to A or B to B) compartmentalized regions. p value was

calculated by two-sided Wilcoxon sum-rank test; whiskers correspond to interquartile range.

F. Network graph show the common and unique enriched terms of A→B or B→A genes,

genes were marked “up” or ”down based on log2FC(HPC/Adherent).

**Supplemental Tables**

Supplemental Table S1. Gene set enrichment analysis results comparison stromal and endothelial cells based on single cell RNA expression.

Supplemental Table S2. Gene set enrichment analysis results comparison endothelial and HE cells based on single cell RNA expression.

Supplemental Table S3. Gene set enrichment analysis results comparison HE cells and HPCs based on single cell RNA expression.

Supplemental Table S4. Changes in transcription factor binding site accessibility based on single cell ATAC sequencing in stromal vs endothelial cells.

Supplemental Table S5. Changes in transcription factor binding site accessibility based on single cell ATAC sequencing in endothelial vs HE cells.

Supplemental Table S6. Changes in transcription factor binding site accessibility based on single cell ATAC sequencing in HE cells vs HPCs.

Supplemental Table S7. Gene ontology analysis identifying pathways based on genes that go from ‘open’ to ‘closed’ based on chromatin accessibility in adherent (stromal/endothelial cell types) vs non-adherent (HPCs) cells.

Supplemental Table S8. Gene ontology analysis identifying pathways based on genes that go from ‘closed’ to ‘open’ based on chromatin accessibility in adherent (stromal/endothelial cell types) vs non-adherent (HPCs) cells.

Supplemental Table S9. Ligand-receptor interaction pairs identified in CellChat analysis of our single cell analysis. All cell types were included in this analysis. Ligands were expressed or secreted in the ‘source’ cell and received by receptors on the surface of ‘target’ cells.

Supplemental Table S10. Antibodies used in this study.

Supplemental Table S11. Summary GWAS statistics used in this study.
