## Supplemental Methods for "Multimodal analysis of *in vitro* hematopoiesis reveals blood cell-specific genetic impacts on complex disease traits"

Christopher S Thom

10-052 Colket Translational Research Building

3501 Civic Center Blvd

Philadelphia, PA 19104

Struan FA Grant

1102D Abramson Research Center

3615 Civic Center Boulevard

Philadelphia, PA 19104

**Supplemental Methods**

**Supplemental Methods**

**Description of *in vitro* hematopoiesis model**

We used a definitive hematopoiesis model that has been previously validated in functional assays^13,20,44,67^. Briefly, iPSCs derived from an anonymous healthy donor were obtained from the CHOP Human Pluripotent Stem Cell Core Facility. These were cultured and dissociated into embryoid bodies (EBs), which were cultured in cytokine mixes that sequentially facilitate production of mesoderm and endothelial cell types. We isolated CD34^+^ endothelial cells using bead selection (StemCell Technologies) and plated aggregates on Matrigel. Subsequent culture in hematopoietic cytokines produces hemogenic endothelial cells and HPCs after 7-9 days. Flow cytometry was performed at serial time points during differentiation on a Cytoflex LX instrument (Beckman Coulter) and analyzed using FlowJo software (BD Biosciences). Cell cultures were imaged on an EVOS imaging system (ThermoFisher Scientific). Antibodies used for these studies are listed in **Supplemental Table S10**.

**Functional hematopoietic assays**

We performed Methocult and Megacult assays according to manufacturer instructions (Stem Cell Technologies). Colonies were quantified and imaged on an Olympus IX70 microscope.

**Cell clustering and annotation**

Single cells were clustered using dimension reduction projections based on RNA expression and the "FindClusters" function. Clusters were annotated based on differentially expressed genes (DEGs) identified using the "FindMarkers" function with default parameters (Seurat v5), manual curation based on literature review, as well as comparison and input from the SingleR and Azimuth data sets.

**Library preparation and Single-cell RNA and ATAC sequencing**

Single-nucleus RNA and ATAC sequencing was performed using the 10x Genomics Chromium Next GEM Single Cell Multiome ATAC + Gene Expression platform according to the manufacturer’s protocol. ATAC and gene expression libraries were generated and sequenced on an Illumina NovaSeqTM 6000 using SBS Cartridge. Library quality was assessed using the Agilent 4200 TapeStation, and final quantification was performed by qPCR (ABI Applied Biosystems ViiA 7). The ATAC library was normalized to 350 pM (1.75 nM, 100 µL) and sequenced on a NovaSeq 6000 SP v1.5 flow cell with a paired-end, dual-indexing configuration (50 × 8 × 24 × 49 cycles) at a depth of 25,000 read pairs per nucleus. The gene expression library was similarly normalized and sequenced on a NovaSeq 6000 S2 v1.5 flow cell using a paired-end, dual-indexing strategy (28 × 10 × 10 × 90 cycles) with a target depth of 20,000 read pairs per nucleus. Data processing was conducted using Cell Ranger 7.1.0 (10x Genomics), with sequence reads aligned to the GRCh38 reference genome for transcript quantification and downstream analysis. The raw output data were processed with the Seurat package (https://satijalab.org/seurat/) in R software (version 4.2.3) for each individual sample. Mitochondrial gene expression was assessed using PercentageFeatureSet (pattern = "^MT-") to identify low-quality cells. ATAC-seq peaks were filtered to retain only those from standard chromosomes (1–22, X, Y) using standardChromosomes(), and peaks detected in fewer than 10 cells were excluded (min.cells = 10). Highly variable ATAC features were selected using FindTopFeatures() with a q0 cutoff.

**Data integration and the dimensionality reduction**

The SCTransform function was applied to normalize the RNA-seq data. Following normalization, Principal Component Analysis (PCA) was performed to reduce the dimensionality of the RNA data. To facilitate further dimensionality reduction and visualization, Uniform Manifold Approximation and Projection (UMAP) was applied to the first 50 principal components from the PCA analysis, generating a 2D UMAP plot for visual representation.

For ATAC-seq data, normalization was carried out using Term Frequency-Inverse Document Frequency (TF-IDF). The FindTopFeatures function was employed to identify the most variable and statistically significant features (genes or peaks) for downstream analysis, with the min.cutoff = 'q0' parameter used to exclude features that failed to meet the quality threshold. Singular Value Decomposition (SVD) was applied as a dimensionality reduction method for the ATAC data. UMAP was subsequently applied to the reduced ATAC data, utilizing the Latent Semantic Indexing (LSI) reduction for input and considering dimensions 2 through 50.

To integrate the RNA and ATAC data, the FindMultiModalNeighbors function was used to construct a combined nearest-neighbor graph, incorporating both the RNA (PCA) and ATAC (LSI) dimensions.

**Trajectory analysis of single cells**

Single-cell pseudotime trajectories were constructed using the Monocle3 package (v2.8.0) in R. The CreateDimReducObject function was utilized to incorporate the UMAP embeddings from the "umap.rna" reduction of the HPC_HE_seurat object, generating a new UMAP object within the Seurat object. This newly created UMAP was then designated as the default for subsequent visualizations. A subset of the Seurat object, consisting of Stromal, Endothelial (Endo), and Hemogenic Endothelial (HE) cells, was converted into a cell_data_set object for trajectory analysis with Monocle. Stromal cells were designated as the root for pseudotime analysis. Monocle performed graph-based differential expression analysis to identify differentially expressed genes (DEGs) by examining gene relationships along the principal trajectory. DEGs were selected based on a q-value threshold of < 0.05, and a plot was generated to illustrate their expression patterns along the pseudotime trajectory.

To further validate the Monocle 3 trajectory analysis, the Slingshot algorithm, a widely used tool for bifurcation trajectory inference, was applied to reanalyze the cellular trajectory of Stromal, Endo, HPC, and HE cells. The getLineages function was employed using UMAP (or PCA) embeddings (dimred) and cluster labels (clustering_factor) to infer the cellular lineages. The SlingMST function was utilized to extract the Minimum Spanning Tree (MST), which represents the shortest path connecting cells along the inferred trajectory. Additionally, the SlingCurves function was used to delineate the Slingshot curves, which represent the trajectories of cells along the distinct lineages.

**Cell-cell interaction analysis**

CellChat (v2.1.1) was employed with default settings to assess intercellular interactions [ref]. The analysis of cell communication was conducted at the cell type level, as specified, with figures and raw output presented accordingly, unless stated otherwise. The “TriMean” method was specified as the argument for the computeCommunProb function. Interactions were examined based on available annotations, specifically for “Secreted Signaling.” Statistical estimates for signaling pathways were directly derived from the CellChat output, which reflects the results of permutation testing, where cell group labels were randomly permuted, and the communication probability recalculated (n = 100 permutations) [ref]. Graphical outputs were generated using CellChat functions.

**Hi-C library processing**

Our workflow follows the pipeline as recently described^1^. Paired-end reads from each replicate were pre-processed using the HICUP pipeline (v0.7.4)^2^, aligned by bowtie2 with hg38 as the reference genome. Each Hi-C library replicate was sequenced to over 3.3 billion reads, and 1.8 billion reads passing quality-control metrics per library were used to construct 3D chromatin maps. The alignments files were parsed to *pairtools* (v0.3.0) to process and *pairix* (v0.3.7) to index and compress, then converted to Hi-C matrix binary format .*cool* by cooler v0.8.11 at multiple resolutions (500bp, 1, 2, 4, 10, 40, 500kbp and 1Mbp) and normalized with ICE method^3^. We then used 40 kb resolution matrices for compartment analysis, 10 kb for TAD analysis, and 1,2 and 4 kb for chromatin loop detection.

We used HiCRep (v1.12.2)^4^ to measure reproducibility across replicates of Hi-C libraries using normalized Hi-C contact matrices at 10 kb resolution, which generates a smoothed contact matrix, and stratifies the matrix by the distance between the interacting regions of chromatin, from which a stratum-adjusted correlation coefficient (SCC) is defined.

**AB Compartment**

We used cooltools (v0.5.1) to determine eigenvectors from an ICE balanced Hi-C matrix with 40kb resolution, the first principal components were used with GC% of genome region as reference track to determine the sign.

**Hierarchical TADs**

We determined hierarchical TADs by directional index using software HiTAD (v0.4.2)^5^ on bireplicated 10kb binned matrix of HE and HPC-Pro. The resulted TAD regions from HiTAD and the original contact matrices were used as input for TADCompare (v1.18.0)^6^ to produce individual boundary scores in HE and HPC, and a classification of the type of differential change observed.

**Chromatin loop calling**

The matrices from different replicates were merged at each resolution using *cooler*. Mustache (v1.0.1)^7^ and Fit-Hi-C2 (v2.0.7)^8^ were used to call significant intra-chromosomal interaction loops from merged replicates matrices at three resolutions 1kb, 2kb, and 4kb, with significance threshold at q-value < 0.1 and FDR < 1 × 10^−6^, respectively. The identified interaction loops were merged between both tools at each resolution. Lastly, interaction loops from all three resolutions were merged with preference for smaller resolution if overlapped.

Quantitative loop differential analysis across cell types was performed on fast lasso normalized interaction frequency (IF) by multiCompareHiC (v1.8.0)^9^ for each chromosome at resolution 1, 2 and 4kb independently. The contacts with zero interaction frequency (IF) among more than 80% of the samples and average IF less than 5 were excluded from differential analysis. The QLF test based on a generalized linear model was performed in cell type-pairwise comparisons, and p values were corrected with FDR. The final differential loops were identified by overlapping differential IF contacts with consensus interaction loops.

**Enrichment analysis**

We performed enrichment analysis using pathfindR (v 2.4.1)^10^ with log2FC and FDR-adjust p-values from differential expression (DE) testing based on the non-parametric Wilcoxon rank sum test.

**Multiome Analysis and eGRN Inference**

Enhancer-mediated gene regulatory networks (eGRNs) were inferred from multiome data using SCENIC+ (v 0.1a1)^11^. The analysis input consisted of the gene expression matrix, imputed accessibility data from pycisTopic (v2.0a0), and TF cistromes previously identified by motif enrichment using pycisTarget (v 1.0a2).

To derive a set of consensus peaks from 10 individual peak sets of each cell type, we used the iterative overlap peak merging procedure as implemented in pycisTopic. First, each summit is extended a ‘peak_half_width’ (by default, 250 bp) in each direction and then we iteratively filtered out less significant peaks that overlap with a more significant one. During this procedure, peaks are merged and, depending on the number of peaks included into them, different processes will happen: (1) 1 peak: the original peak will be retained; (2) 2 peaks: the original peak region with the highest score will be retained; and (3) 3 or more peaks: the original region with the most significant score will be taken, and all of the original peak regions in this merged peak region that overlap with the significant peak region will be removed. This resulted in 336,932 regions. We further filtered the dataset on the basis of the snATAC–seq quality as well, retaining cells with at least 1,000 fragments, FRiP > 0.4 and TSS enrichment > 7, resulting in 23,420 high-quality cells. Topic modelling was performed using Mallet (v.2.0), using 500 iterations and models with 2 topics and from 5 to 100 by an increase of 5. Additional models between 25 and 55 (by an increase of 1) were added, as we observed that the best model should be on that area based on the model selection metrics, and we selected a model with 43 topics. Batch effects between samples were corrected using harmonypy (v.0.0.6) on the scaled topic distributions, and Leiden clustering with a resolution of 0.6 resulted in 11 clusters, corresponding to 10 cell types based on previous labelling. Drop-out imputation was performed by multiplying the region-topic and topic-cell probabilities. The imputed accessibility matrix was multiplied by 10^6^.

SCENIC+ was run with default parameters on the complete multiome dataset. Region-to-gene relationships were determined using gradient boosting machine regression. A search space was defined as a minimum of 1 kb upstream of the TSS or downstream of the gene end, extending to a maximum of either the closest gene boundary or 1 Mb. TF-to-gene relationships were also calculated using gradient boosting. eRegulons were built via a GSEA-based method, employing binarization of region-to-gene scores using multiple thresholds (85th, 90th, 95th quantiles) and selecting the top regions per gene (5, 10, 15), alongside the BASC method^12^.

**Definition of cis-Regulatory Elements (cREs)**

We intersected ATAC-seq open chromatin regions (OCRs) of each cell type with chromatin conformation capture data determined by Hi-C of the same cell type group, and with promoters (-1,500/+500bp of TSS) defined by GENCODE v40.

**Reformatting GWAS summary statistics**

**Supplemental Table S11** lists the studies from which we drew GWAS summary statistics for each trait.

**LDSC**

This package recommends reformatting the summary statistics files using *munge_sumstats.py*. The table shows the number of total variants reported for each trait from the original studies, and the numbers after being filtered by *munge_sumstat.py*. This step also produced basic metadata about the summary statistics, such as mean chi^2^, max chi^2^, lambda GC, and mean value of the signed summary statistic column (beta, odd ratio - OR). Due to the inability of ***LDSC*** to produce meaningful heritability *h^2^* for traits with low numbers of variants, we applied *--merge-alleles* with the list of HapMap3 variants that LDSC used to estimate LD Scores (<https://data.broadinstitute.org/alkesgroup/LDSCORE/w_hm3.snplist.bz2>). This standardized all the GWAS sumstats files to the 1,217,311 variants from HapMap, discarded all the unmatched, and imputed the missing variants into the sumstats with *NULL* beta value.

**Partitioned heritability for cell type specific annotations with each trait**

We applied S-LDSC (stratified linkage disequilibrium score regression) using LDSC v.1.0.1 to quantify to contribution to SNP-based effect-sizes and heritability to 19 complex diseases from each cell type’s cREs. Each set of input regions from each cell type was used to create the annotation, which in turn was used to compute annotation-specific LD scores for each cell types regions of interest set. These annotation-specific LD scores were used with 63 categories of the full baseline model. (v2.2) as control. Two statistics we used to assess how effectively our annotations capture causal variation are heritability enrichment and standardized effect size (τ*), as previously defined^13^. When conditioning two correlated annotations in a joint S-LDSC model, they may show similar enrichments, but the τ* for the annotation with a higher true causal variant membership will be larger and more positive.

**Genetic loci included in variant-to-genes mapping**

For “Heights” and “T1D” whose studies performed fine-mapping analysis, we used the significant variants within their provided credible sets. For the other traits, we leveraged their reported variants from associated loci which reached genome-wide significance within original studies. Proxies for each lead variant were queried using TopLD^14^ and LDlinkR tool^15^ with the GRCh38 Genome assembly, 1000 Genomes phase 3 v5 variant set, European population, and LD threshold of r^2^≥0.8 (results in **Supplemental Table S11**, column “N proxies”).
